# Characterization of an evolutionarily distinct bacterial ceramide kinase from *Caulobacter crescentus*

**DOI:** 10.1101/2023.05.01.538943

**Authors:** Tanisha Dhakephalkar, Geordan Stukey, Ziqiang Guan, George M. Carman, Eric A. Klein

## Abstract

A common feature among nearly all Gram-negative bacteria is the requirement for lipopolysaccharide (LPS) in the outer leaflet of the outer membrane. LPS provides structural integrity to the bacterial membrane which aids bacteria in maintaining their shape and acts as a barrier from environmental stress and harmful substances such as detergents and antibiotics. Recent work has demonstrated that *Caulobacter crescentus* can survive without LPS due to the presence of the anionic sphingolipid ceramide-phosphoglycerate. Based on genetic evidence, we predicted that protein CpgB functions as a ceramide kinase and performs the first step in generating the phosphoglycerate head group. Here, we characterized the kinase activity of recombinantly expressed CpgB and demonstrated that it can phosphorylate ceramide to form ceramide 1-phosphate. The pH optimum for CpgB was 7.5, and the enzyme required Mg^2+^ as a cofactor. Mn^2+^, but not other divalent cations, could substitute for Mg^2+^. Under these conditions, the enzyme exhibited typical Michaelis-Menten kinetics with respect to NBD-C6-ceramide (K_m,app_=19.2 ± 5.5 μM; V_max,app_=2586.29 ± 231.99 pmol/min/mg enzyme) and ATP (K_m,app_=0.29 ± 0.07 mM; V_max,app_=10067.57 ± 996.85 pmol/min/mg enzyme). Phylogenetic analysis of CpgB revealed that CpgB belongs to a new class of ceramide kinases which is distinct from its eukaryotic counterpart; furthermore, the pharmacological inhibitor of human ceramide kinase (NVP-231) had no effect on CpgB. The characterization of a new bacterial ceramide kinase opens avenues for understanding the structure and function of the various microbial phosphorylated sphingolipids.

## Introduction

Gram-negative bacteria have a three-layered cell envelope composed of the inner membrane, a thin layer of peptidoglycan-cell wall and an outer membrane. A key component of the outer membrane is lipopolysaccharide (LPS) (1). LPS is an essential molecule in nearly all Gram-negative species due to its roles in barrier formation and membrane integrity (2). While the general structure of LPS is well conserved, there is considerable variation between and within species (3). LPS can be divided into three structural domains: 1) lipid A, a membrane anchored multi-acylated oligosaccharide, 2) the core oligosaccharide, often containing 3-deoxy-d-manno-oct-2-ulosonic acid (Kdo), which is generally conserved within a species, and 3) the polysaccharide O-antigen, which is highly variable, even among strains of the same species. In many organisms, like *Escherichia coli*, the lipid A portion of LPS is negatively charged due to the presence of phosphate groups on the glucosamine disaccharide (3). These phosphates are the binding sites for cationic antimicrobial peptides (CAMPs) like polymyxins (4,5). While LPS is generally considered to be essential, LPS-null mutants of several Gram-negative organisms have been isolated including *Acinetobacter baumannii* (6), *Moraxella catarrhalis* (7), *Neisseria meningitidis* (8), and *Caulobacter crescentus* (9). The ability of *C. crescentus* to survive in the absence of LPS is, in part, due to the presence of the anionic sphingolipid ceramide-phosphoglycerate (CPG), as sphingolipid synthesis becomes essential in the LPS-null mutant (9). In contrast to *E. coli*, the mature lipid A molecule in *C. crescentus* is not phosphorylated; instead, the phosphate groups are hypothesized to be removed by the phosphatase CtpA (9) and replaced with galactopyranuronic acid (10). Whereas polymyxin antibiotics target the phosphorylated lipid A in *E. coli*, antibiotic sensitivity assays demonstrated that CAMPs kill *C. crescentus* by interacting with the anionic CPG lipids. Synthesis of the CPG headgroup is sequentially catalysed by the three proteins CpgABC (CCNA_01217-01219) (9) (Figure 1A). Deletion of *cpgB* (*ccna_01218*) results in the loss of ceramide 1-phosphate (C1P) which is consistent with its annotation as a putative lipid kinase (9).

**Figure 1:**
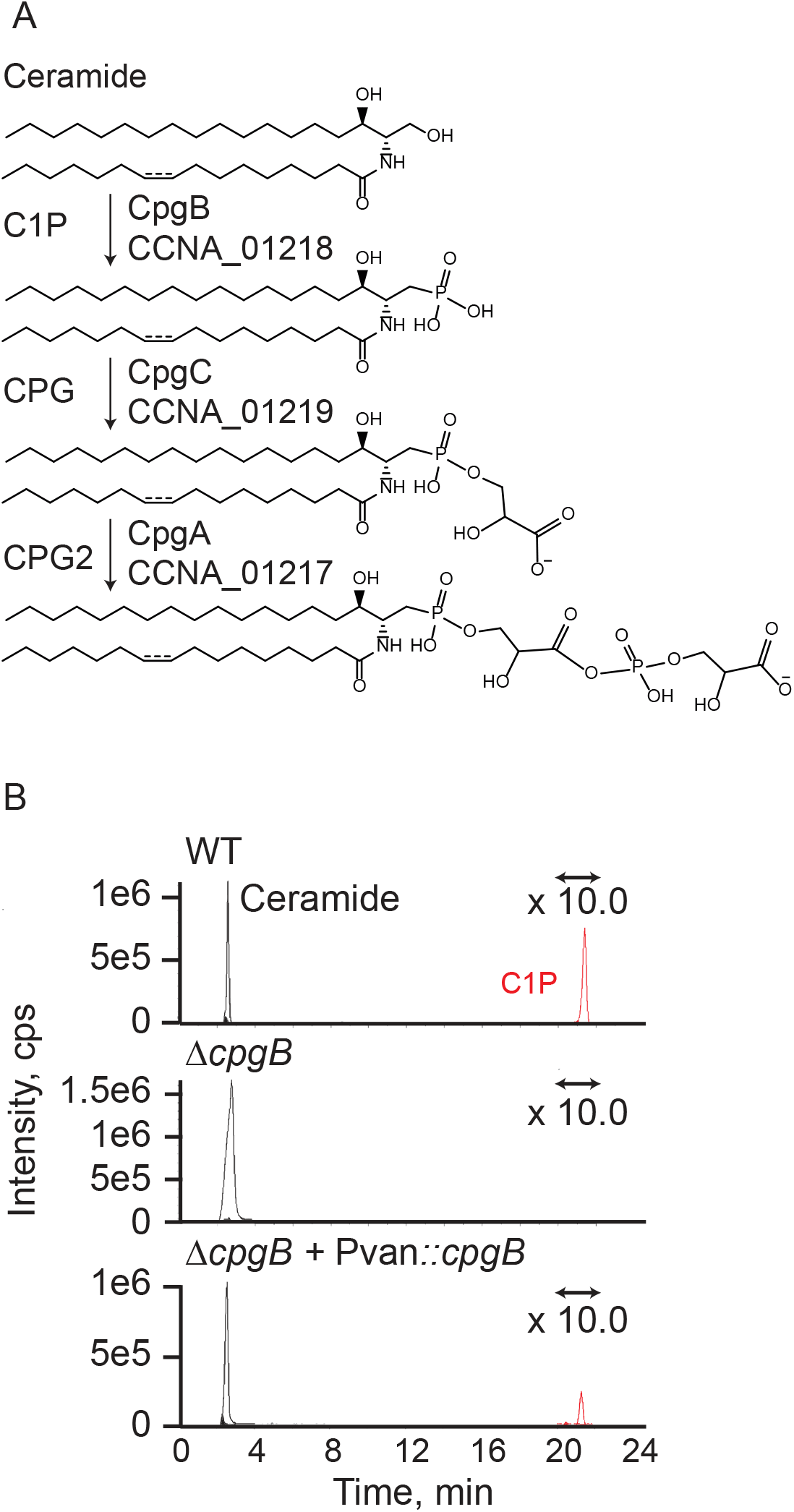
Identification of CpgB as a putative ceramide kinase. (A) Previous genetic analysis of the *cpgABC* genes led to a proposed mechanism for ceramide-phosphoglycerate (CPG) synthesis (9). (B) Extracted-ion chromatograms show the presence or absence of ceramide and C1P. Total lipids were extracted from the indicated strains and analyzed by normal phase LC/ESI-MS in the negative ion mode. The signal for the C1P peak was magnified 10-fold since this lipid is only a minor component of the *C. crescentus* lipidome.

C1P has important physiological roles in eukaryotes including mast cell activation, phagocytosis, cellular proliferation, and survival (reviewed in (11)). Human ceramide kinase (hCERK) uses ceramide and ATP as substrates to produce C1P (12). The CERK enzyme is part of a larger family of lipid kinases including sphingosine kinase and diacylglycerol kinase. A bacterial dihydrosphingosine kinase has recently been identified in *Porphyromonas gingivalis* (13); however, to our knowledge, this is the first described bacterial CERK enzyme. In this study we used purified *C. crescentus* CpgB to characterize its ceramide kinase activity. Phylogenetic analysis comparing various lipid kinases suggests that bacterial CERK enzymes form a distinct clade from their eukaryotic counterparts.

## Results

### CCNA_01218 is a bacterial ceramide kinase

Most Gram-negative bacteria require LPS in the outer membrane for survival. A recently isolated mutant of *C. crescentus* is capable of surviving without LPS, largely due to the presence of the anionic sphingolipid CPG (9). Genetic analysis identified 3 genes (*ccna_01217-01219*) that were required for synthesizing the phosphoglycerate headgroup. CCNA_01218 (CpgB) is annotated as a lipid kinase-related protein and deletion of *cpgB* resulted in a loss of C1P (Figure 1B) (9), consistent with *cpgB* encoding a bacterial ceramide kinase. To determine the enzymatic activity of CpgB, we purified the His-tagged recombinant protein from *E. coli* (Figure 2) and performed kinase assays. CpgB could readily phosphorylate C16-ceramide (Figure 3A) as well as a fluorescent NBD-C6 ceramide (Figure 3B). The identity of the phosphorylated NBD-ceramide product was confirmed by mass spectrometry (Figure 3C). Since CpgB has a conserved LCB5 diacylglycerol (DAG) kinase domain, we tested whether CpgB could phosphorylate DAG to produce phosphatidic acid (PA) and found comparable activity (Figure 1A-B). Although CpgB can produce PA *in vitro*, the *C. crescentus* lipidome contains only ∼1% PA (14) and deletion of *cpgB* had no effect on PA levels (Figure 3D). Therefore, we conclude that ceramide is the preferred *in vivo* substrate for CpgB.

**Figure 2:**
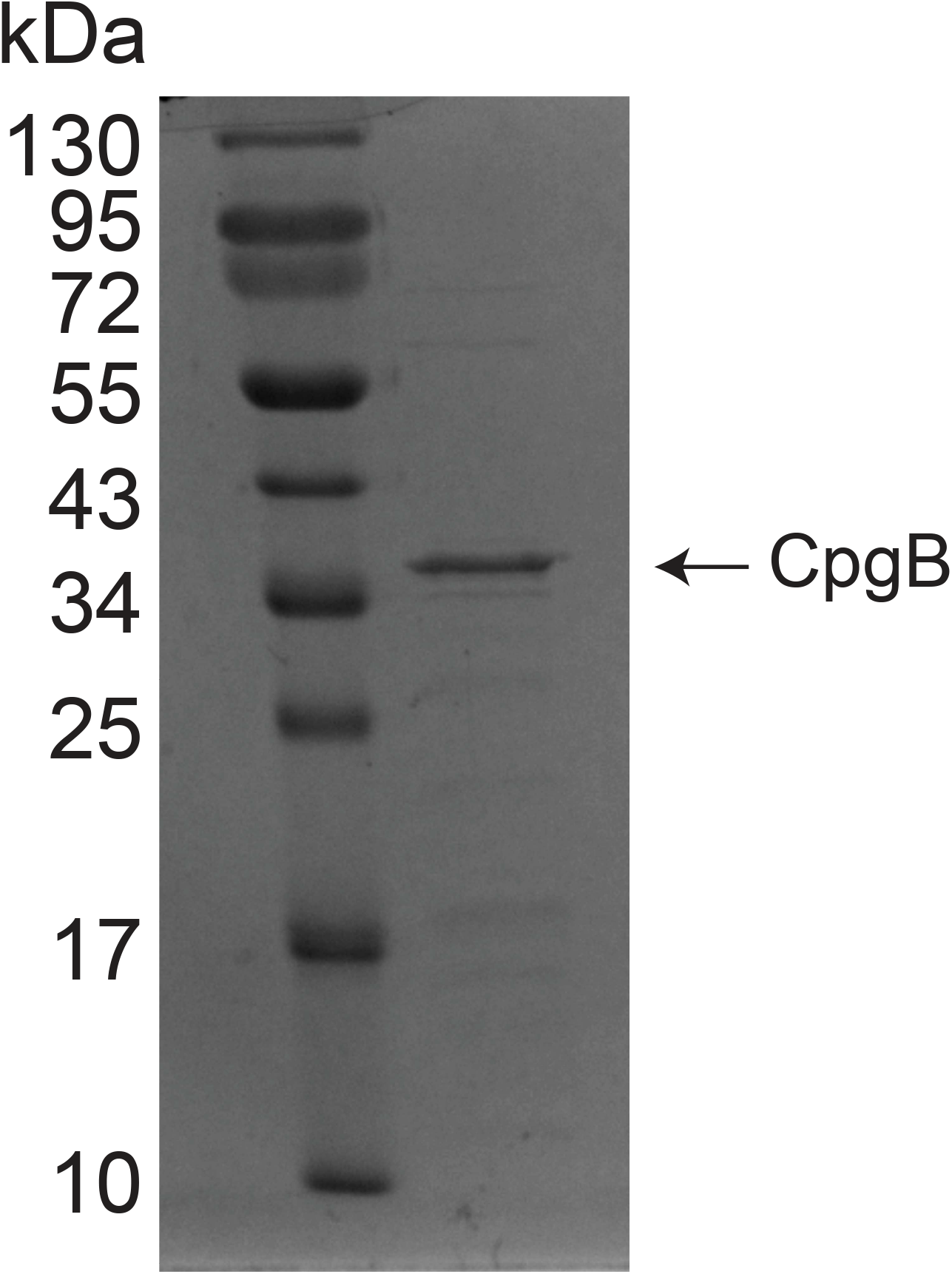
Purification of CpgB. His-tagged CpgB was expressed and purified from *E. coli*. An SDS-PAGE gel of recombinant CpgB was stained with Coomassie Blue to assess protein purity.

**Figure 3:**
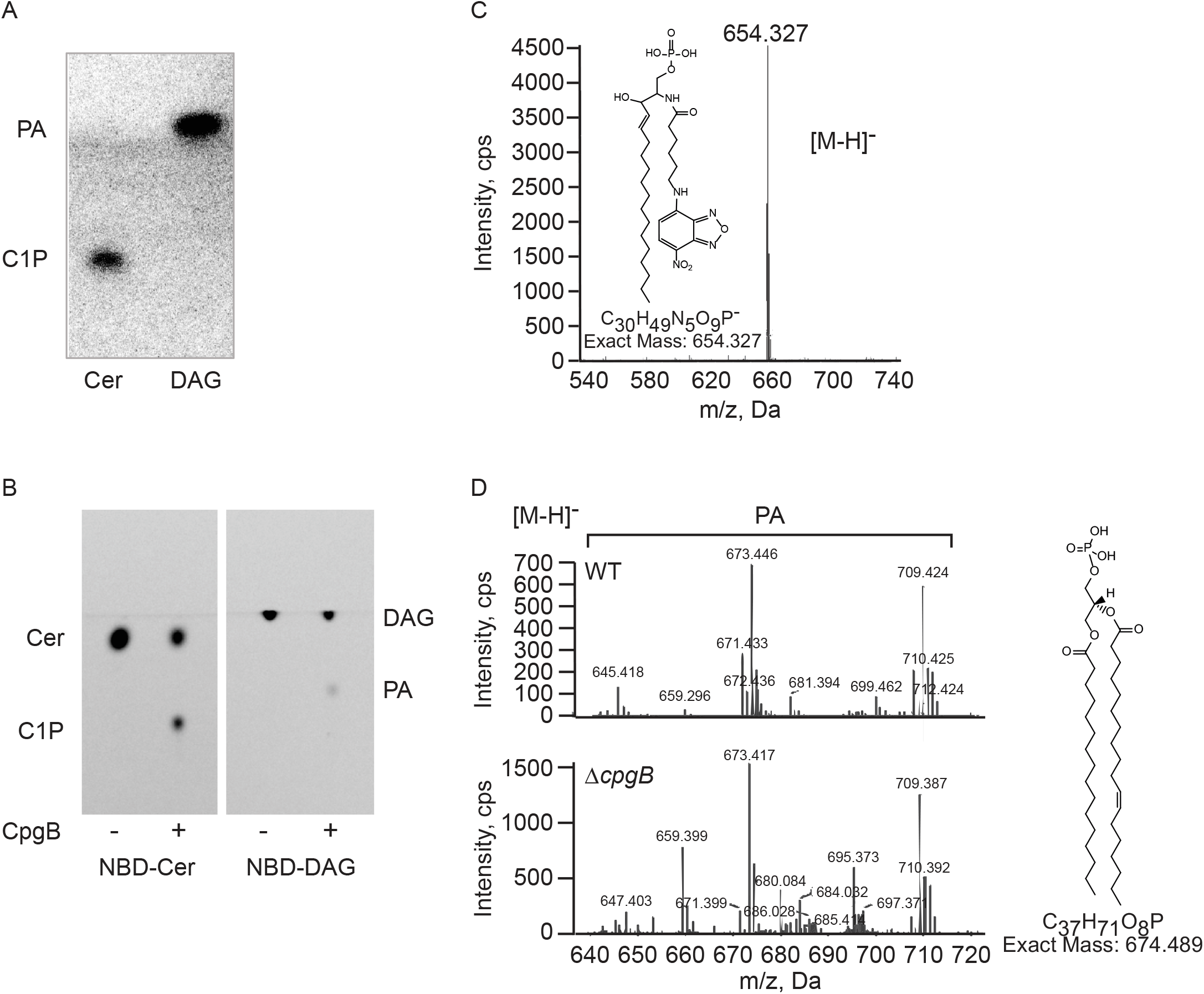
Cpg has ceramide kinase activity. (A) Recombinant CpgB was used to phosphorylate C16-ceramide or DAG. ^32^P incorporation was monitored by TLC and phosphorimaging. (B) The substrate specificity of CpgB was analyzed using fluorescent NBD lipid substrates as indicated. (C) Production of the phosphorylated NBD-ceramide product was confirmed by negative ion ESI/MS analysis. (D) Negative ion ESI/MS analysis of lipid extracts from wild type (WT) and Δ*cpgB* strains shows no difference in phosphatidic acid (PA) levels.

### Influence on pH and divalent cations on CpgB activity

To characterize the requirements for CpgB activity, was measured C1P production over a pH range from 4.5-10; optimal activity was observed at pH 7.5 (Figure 4A). By contrast, hCERK has optimal activity at pH 6.5 (12,15). Since hCERK activity increases strongly in the presence of magnesium or calcium (12), we tested CpgB’s dependence on divalent cations. In the absence of any cations, we did not observe production of C1P (Figure 4B). Both magnesium and manganese strongly increased CpgB activity, with smaller effects observed in the presence of zinc or cobalt (Figure 2C). In contrast to hCERK, calcium did not stimulate CpgB activity (Figure 4C).

**Figure 4:**
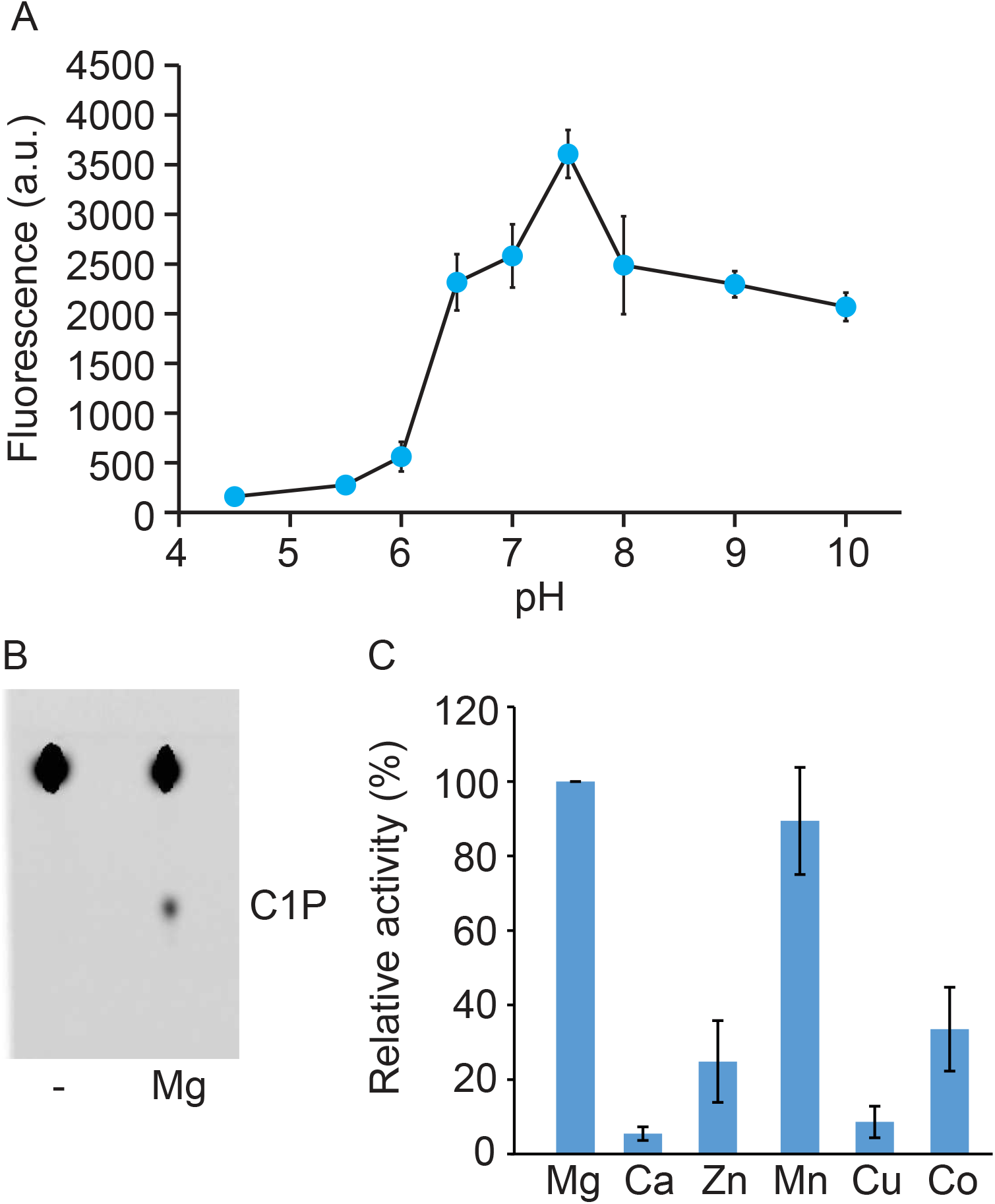
Characterization of CpgB pH and divalent cation requirements. (A) CpgB kinase activity was determined over a range of pH’s using the following buffers: pH 4.5–6 (100 mM citrate), pH 6.5–7.5 (100 mM MOPS), pH 8–9 (100 mM Tris-HCl) and pH 10 (100 mM borate). Activity was quantified using the NBD-ceramide substrate (n=3, error bars are the SD). (B) The CpgB kinase assay was performed in the presence or absence of 15 mM Mg^2+^ using NBD-ceramide. Product formation was analyzed by TLC. (C) The activity of CpgB was determined in the presence of 15 mM of the indicated divalent cations. Activities were normalized to magnesium (n=3; error bars are the SD).

### Determination of CpgB kinetic parameters

Using the NBD-ceramide substrate, we measured C1P production over a two-hour period to identify the linear range of activity for subsequent determinations of enzyme kinetic parameters (Figure 5A); unless otherwise noted, all remaining kinase assays were performed for 30 minutes in the presence of Mg^2+^ at pH 7.4. The enzyme exhibited typical Michaelis-Menten kinetics with respect to NBD-C6-ceramide (K_m,app_=19.2 ± 5.5 μM; V_max,app_=2586.29 ± 231.99 pmol/min/mg enzyme) and ATP (K_m,app_=0.29 ± 0.07 mM; V_max,app_=10067.57 ± 996.85 pmol/min/mg enzyme) (Figures 5B-C). We are reporting apparent K_m_ and V_max_ values since CpgB has two substrates and performs a Bi-Bi reaction; under these conditions the concentration of each substrate affects the apparent kinetic parameters of the other.

**Figure 5:**
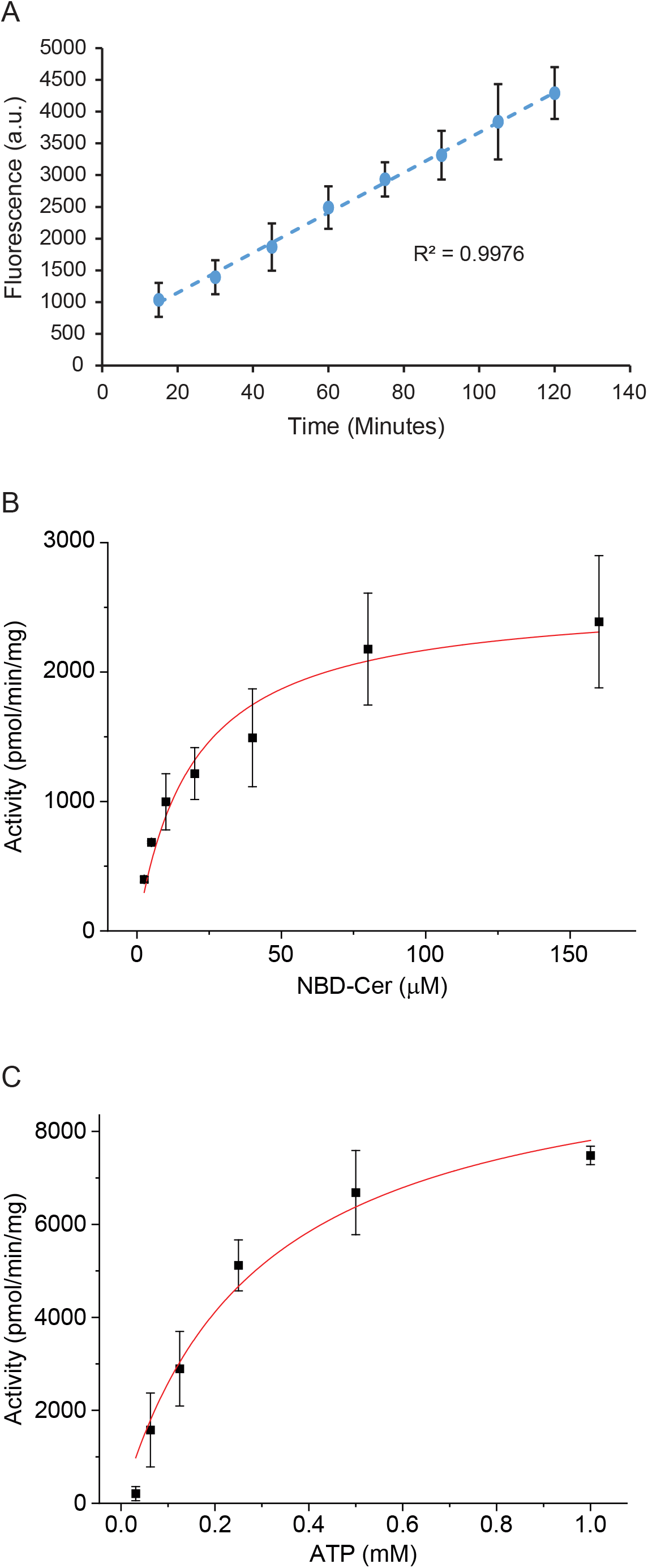
CpgB enzyme kinetics. The kinetic parameters of CpgB were measured using the C6-NBD ceramide substrate. (A) CpgB activity was measured as a function of time (n=3, error bars are SD). (B-C) Michaelis-Menten kinetic parameters were determined for CpgB (n=2, error bars are SD). (B) To determine the K_m,app_ for ceramide, ATP concentration was held constant (1 mM) while NBD-ceramide concentration varied. (C) The K_m,app_ for ATP was determined by holding the NBD-ceramide constant at 160 μM while varying the ATP concentration. K_m,app_ values were 19.2 ± 5.5 μM and 0.29 ± 0.07 mM for NBD-ceramide and ATP, respectively. V_max,app_ values were 2586.29 ± 231.99 pmol/min/mg enzyme and 10067.57 ± 996.85 pmol/min/mg enzyme for NBD-ceramide and ATP, respectively.

### Bacterial and eukaryotic ceramide kinases are phylogenetically distinct enzymes

Given the observed enzymatic differences between hCERK and CpgB, we considered whether these two enzymes are evolutionarily related. Sequence alignment shows limited agreement (12.5% identity and 22.5% similarity); four of the five sphingosine kinase conserved domains show some homology between the eukaryotic and bacterial kinases (Figure 6A) (12). The two kinases also share a conserved GGDG motif which is involved in ATP binding (16). However, the eukaryotic ceramide kinases have an absolutely conserved CxxxCxxC motif that is required for enzyme activity (17) but is absent from CpgB.

**Figure 6:**
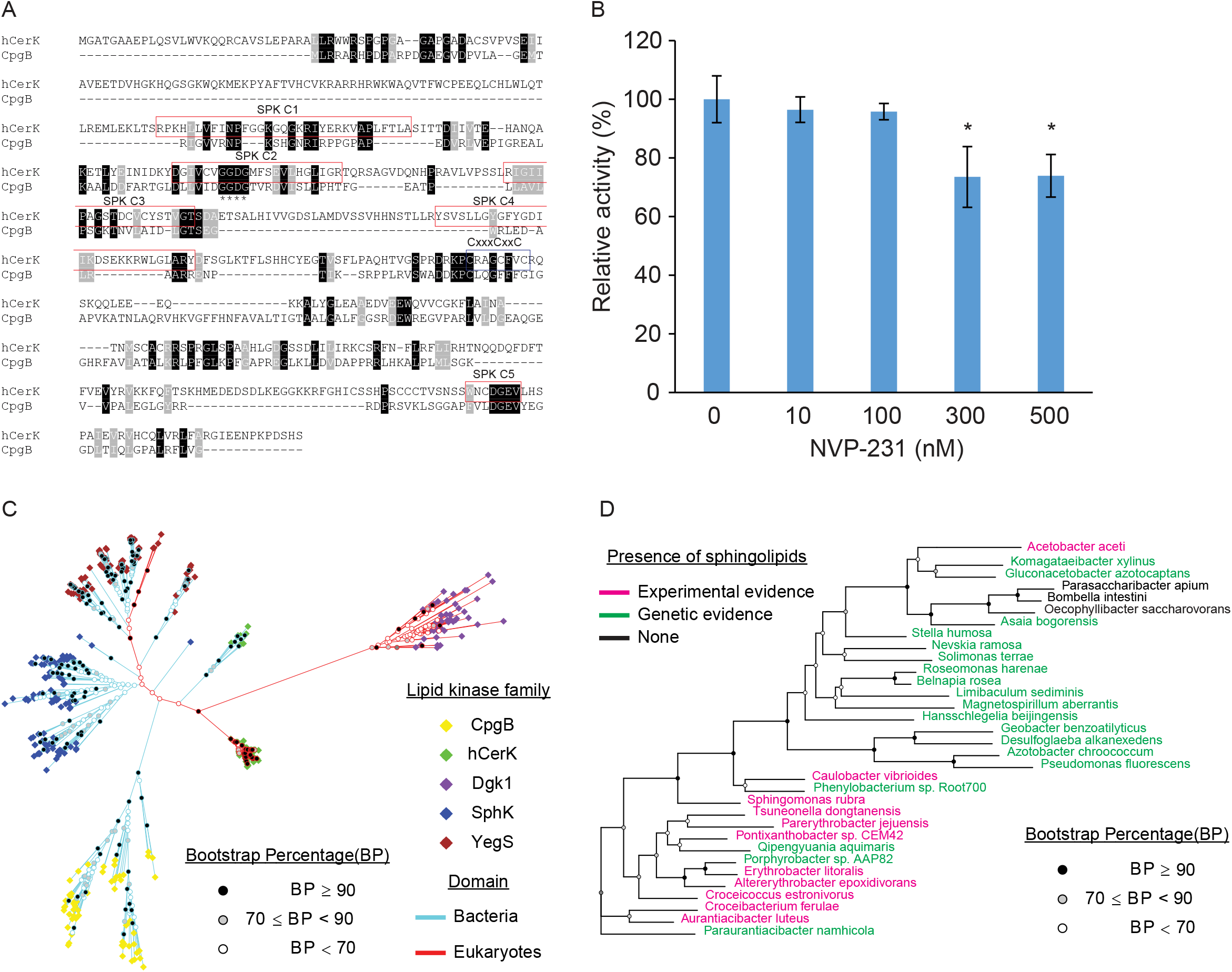
Bacterial CERK is a unique class of lipid kinases. (A) Sequence alignment of CpgB and hCERK shows limited homology. The hCERK sphingosine-kinase domains are indicated by the red boxes. The conserved GGDG motif is indicated with asterisks. hCERK has a conserved CxxxCxxC which is absent in CpgB. (B) CpgB activity was measured in the presence of the hCERK inhibitor NVP-231 using the NBD-ceramide substrate. Activities were normalized to the control sample (n=3; error bars are SD, ANOVA F(4,10)=10.22, P<0.0015; * post-hoc comparisons using Tukey test, P<0.05). (C) Phylogenetic analysis of various lipid kinases was performed using the maximum-likelihood method. The branch-tip color indicates the lipid-kinase family, and the line colors designate proteins of bacterial or eukaryotic origin. (D) The phylogenetic tree of the bacterial CpgB homologues is color coded to indicate which genera have members with either experimental evidence (pink) or genetic evidence (green) of sphingolipid production. Genetic evidence indicates that the genus has members with all three required enzymes for sphingolipid production: Spt, bCerS, and CerR.

To further assess the functional similarity between the ceramide kinases, we treated CpgB with the hCERK inhibitor NVP-231 (18). NVP-231 is a competitive inhibitor of ceramide binding and inhibits 90% of hCERK activity at 100 nM (18). By contrast, 100 nM NVP-231 had no significant effect on CpgB activity (Figure 4B). When the concentration was increased to 300 nM, we observed only a modest 20-25% inhibition (Figure 6B).

Several enzyme families are capable of phosphorylating sphingolipids and DAG. To visualize the similarity of CpgB to these enzymes, we performed a maximum-likelihood phylogenetic analysis and included representative proteins from the following families: hCERK, yeast diacylglycerol kinase Dgk1 (19), bacterial dihydrosphingosine kinase dhSphK1 (13), and bacterial phosphatidylglycerol kinase YegS (20). Each of these enzymes formed a distinct clade despite having overlapping activities (Figure 6C). We did find several cyanobacterial enzymes with homology to hCERK as well as some green algae with homologues of YegS; CpgB homologues were only found in bacterial species. Further analysis of the CpgB-encoding organisms revealed that nearly all genera with the *cpgB* gene either produce or encode the genes required for sphingolipid synthesis (21) (Figure 6D).

## Discussion

Bacterial sphingolipids have a wide range of head groups including sugars (22-24), phosphoglycerol (25), phosphoglycerate (9), and phosphoethanolamine (26). These modifications likely determine the physiological functions of the respective sphingolipids. For example, phosphoglycerol dihydroceramide produced by *P. gingivalis* promotes osteoclastogenesis through its interactions with non-muscle myosin II-A (25). In the case of *C. crescentus*, production of the anionic CPG enables survival in the absences of LPS (9). Genetic analysis using single-gene deletion mutants led to the identification of three enzymes required for CPG synthesis and suggested the first step is catalyzed by CpgB, a putative ceramide kinase. In this report, we used recombinant CpgB expressed and purified from *E. coli* to confirm its ceramide kinase activity and analyze its enzymatic properties.

CpgB differs from the human CERK with regards to divalent cation specificity, susceptibility to the inhibitor NVP-231, and kinetic parameters. For comparison, the K_m,app_’s for CpgB are 19 μM and 0.29 mM for ceramide and ATP, respectively, whereas the reported K_m_’s for hCERK are 187 μM and 32 μM (12).

From a structural perspective, hCERK activity is observed in cellular membrane fractions despite not having any predicted transmembrane domains; one explanation is that the N-terminal PH domain interacts with membrane phosphatidylinositol molecules (12). By contrast, CpgB purifies as a soluble protein without the use of detergents and is predicted to be a cytoplasmic protein (27). Consistent with these biochemical findings, phylogenetic analysis suggests that the bacterial CERK forms a unique subfamily of lipid kinases, distinct from the eukaryotic CERK. Broad conservation of CpgB across many classes of bacteria suggests that phosphorylation may be a common sphingolipid modification.

Until recently, the genes responsible for specific ceramide modifications were unknown. As a result, various studies broadly determined the importance of total sphingolipid production by knocking out the *spt* gene and assessing phenotypes related to survival or virulence (26,28,29). With the discovery of enzymes required for sphingolipid glycosylation, phosphorylation, and other modifications (9,13,22,24), we can now dissect the roles of specific headgroup modifications. The characterization of a new bacterial CERK opens avenues for understanding the structure and function of the various microbial phosphorylated sphingolipids.

## Materials and Methods

### Cloning His-tagged CpgB

The *cpgB* gene was amplified from *C. crescentus* genomic DNA using primers EK1462 (tatattcatATGCTTCGTCGTGCACGCCATCC) and EK1464

(tactgaattcTCATCCGACCAGGAACCGCAAGGC) and ligated into the NdeI/EcoRI site of plasmid pET28a to generate an N-terminal His-tagged fusion. The resulting plasmid was verified by Sanger sequencing and transformed into *E. coli* strain BL21(DE3) for expression and purification.

### Purification of CpgB

A 1L culture of *E. coli* BL21(DE3) cells carrying the pET28a-*cpgB* plasmid was grown in LB broth with 30 μg/mL kanamycin at 37 °C with shaking to an OD_600_ of 0.6. Isopropyl ß-D-1-thiogalactopyranoside (IPTG) was added to a final concentration of 0.5 mM, followed by induction at 16 °C for 18 h. The cells were collected by centrifugation at 10,000 x *g* and resuspended in 12.5 mL of buffer containing 0.5 M sucrose and 10 mM Tris, pH 7.5. Lysozyme was added to a final concentration of 144 μg/mL and the suspension was stirred on ice for 2 min. 12.5 mL of 1.5 mM EDTA was added with stirring for an additional 7 min to induce plasmolysis. The cells were collected by centrifugation at 10,000 x *g* for 10 min and the pellet was resuspended in lysis buffer (20 mM Tris pH 7.5, 0.5 M NaCl, 10 mM imidazole) prior to lysis via 2-3 passages through a French press (20,000 psi). The lysate was centrifuged at 8,000 x *g* for 10 min to remove unbroken cells. His-CpgB was purified using an ÄKTA start FPLC system and a 1 mL HisTrap HP column (Cytiva). After loading, the column was washed with lysis buffer prior to elution via a linear gradient to 1M imidazole. Protein elution was monitored by A_280_ and fractions were collected and analyzed by SDS-PAGE followed by Coomassie blue staining. Fractions containing the purified CpgB were combined and dialyzed into 10 mM Tris, pH 7.2, 0.1 M NaCl, 2 mM EDTA, 1 mM DTT over 48 hr at 4 °C. The dialyzed protein was concentrated using an Amicon Ultra centrifugal filter (10 kDa molecular weight cutoff) (Millipore Sigma). The protein concentration was determined using the BCA Protein Assay Kit (Pierce).

### CpgB kinase assay using C16-ceramide

CpgB kinase activity was measured for 30 min at 30 °C as described previously for *E. coli* diacylglycerol kinase (30). The reaction mixture contained 50 mM imidazole-HCl, pH 6.6, 50 mM octyl-β-D-glucopyranoside, 50 mM NaCl, 12.5 mM MgCl_2_, 1 mM EGTA, 10 mM β-mercaptoethanol, 1 mM cardiolipin, 0.1 mM ATP (1000 cpm/pmol), and 0.8 mM ceramide or DAG in a total volume of 20 μl. The radioactive products (PA or C1P) are chloroform soluble and were separated from the remaining radioactive substrate by a non-acidic chloroform/methanol/MgCl_2_ (1 M) phase separation. The chloroform soluble products were separated by TLC using a chloroform/methanol/water (65:25:4, v/v) solvent system and visualized by phosphorimaging.

### NBD-ceramide kinase assay

The NBD-ceramide kinase assays were carried out largely as previously described for human ceramide kinase (12,15). Briefly, the reaction was carried out in a buffer containing: 20 mM HEPES (pH 7.4), 10 mM KCl, 15 mM MgCl_2_, 10% glycerol, 1 mM DTT, 1 mM ATP, 0.2 mg/mL fatty acid-free BSA, and 10 μM C6-NBD ceramide (added from a 10 mM ethanol stock) (Thermo Fisher Scientific). The reaction was started by adding 0.025 μg/μL of the CpgB enzyme. Tubes were incubated in the dark at 30 °C for the indicated times. After the incubation, 1 μL of the reaction mixture was spotted onto Silica Gel 60 TLC plates. The spots were resolved in a solvent system containing butanol/acetic acid/water (3:1:1, v/v). The dried TLC plates were visualized using the GFP filter set on a Bio-Rad ChemiDoc. To test the specificity of CpgB, we performed the reaction under identical conditions using 1-NBD-decanoyl-2-decanoyl-sn-Gly (NBD-DAG) (Cayman Chemical) as the substrate. Inhibition of CpgB activity was performed by adding the indicated concentrations of NVP-231 (Cayman Chemical) to the reaction prior to addition of the enzyme.

### Lipidomic profiling and confirmation of ceramide-phosphate production by LC/MS/MS

Lipids were extracted from bacterial cells or the NBD-ceramide CpgB reaction using the method of Bligh and Dyer with minor modifications (31). The lipid extracts were analyzed by normal phase LC/MS/MS in the negative ion mode as previously described (32,33).

### Kinetic analysis of CpgB

To determine the kinetic constants for CpgB, activity assays were performed for 30 min as described above while varying substrate concentrations. To determine the K_m,app_ for ceramide, ATP concentration was held constant (1 mM) while NBD-ceramide concentration ranged from 0.625-160 μM. The K_m,app_ for ATP was determined by holding the NBD-ceramide constant at 160 μM, while varying the ATP concentration from 0.031-1 mM. Product formation was measured from the fluorescent images using ImageJ (34) and quantified using a standard curve of NBD-ceramide spotted onto the TLC plates. The enzyme activity was fit to the Michaelis-Menten equation using OriginPro (OriginLab).

### Assessing the pH optimum and the requirement for divalent cations

To test the effect of pH on CpgB activity, a standard reaction mix was made containing 10 mM KCl, 15 mM MgCl_2_, 10% glycerol, 1 mM DTT, 1 mM ATP, 0.2 mg/mL fatty acid-free BSA, and 10 μM C6-NBD ceramide. The pH was controlled by adding the following buffers: pH 4.5–6 (100 mM citrate), pH 6.5–7.5 (100 mM MOPS), pH 8–9 (100 mM Tris-HCl), and pH 10 (100 mM borate). The reactions were started with the addition of 0.025 μg/μL of CpgB and allowed to run for 30 min. Phosphorylated product was quantified as above. The efficacy of various divalent cations was tested by replacing the MgCl_2_ with 15 mM CaCl_2_, ZnCl_2_, MnCl_2_, CuCl_2_, or CoCl_2_ and determining CpgB activity at pH 7.4 as described above.

### Phylogenetic analysis of lipid kinase enzymes

Using CCNA_01218 (CpgB; Accession YP_002516591.3) protein as a query, we performed BLASTP searches to find related proteins in the NCBI database (35). The top hits were all from species closely related to *C. crescentus*, so we repeated the search excluding Alphaproteobacteria to get a wider range of organisms. Candidate hits were chosen using an E-value cutoff of 1E-20 and we manually curated the list to select the top ∼60 hits from different genera. A similar method was used to find homologues of hCERK (Accession NP_073603.2), *Porphyromonas gingivalis* dihydrosphingosine kinase (Accession AAQ66413), *E. coli* YegS (Accession NP_416590), and *Saccharomyces cerevisiae* Dgk1 (Accession QHB11896.1). A total of 397 protein sequences were aligned using MUSCLE aligner (36). Phylogenetic trees were prepared using RAxML (Randomized Axelerated Maximum Likelihood version 8.2.12) (37) with 100 bootstraps and a maximum-likelihood search. RAxML was run on the CIPRES Portal at the San Diego Supercomputer Center (38). Phylogenetic trees were visualized in R using the packages ggtree (39), ape (40), treeio (41), and ggplot2 (42). To determine which *cpgB* encoding bacterial genera produce sphingolipids, we performed a literature search as well as used the Riken JCM catalogue (https://jcm.brc.riken.jp/en/). For genera with no experimental evidence of sphingolipids, we used BLASTP to determine whether these genera encode all three key enzymes for sphingolipid synthesis: Spt (Accession A0A0H3C7E9.1), bCerS (Accession A0A0H3C8X0.1), and CerR (Accession A0A0H3C8X7.1).

## Data availability

All of the data for this work is contained within the manuscript.

### Acknowledgements

We thank Valerie Carabetta and Olaitan Akintunde (Cooper Medical School of Rowan University) for their assistance with protein purification.

## References

1. Sutcliffe, I. C. (2010) A phylum level perspective on bacterial cell envelope architecture. Trends Microbiol 18, 464–470

2. Bertani, B., and Ruiz, N. (2018) Function and biogenesis of lipopolysaccharides. EcoSal Plus 8

3. Whitfield, C., and Trent, M. S. (2014) Biosynthesis and export of bacterial lipopolysaccharides. Annu Rev Biochem 83, 99–128

4. Manioglu, S., Modaresi, S. M., Ritzmann, N., Thoma, J., Overall, S. A., Harms, A., Upert, G., Luther, A., Barnes, A. B., Obrecht, D., Muller, D. J., and Hiller, S. (2022) Antibiotic polymyxin arranges lipopolysaccharide into crystalline structures to solidify the bacterial membrane. Nat Commun 13, 6195

5. Morrison, D. C., and Jacobs, D. M. (1976) Binding of polymyxin B to the lipid A portion of bacterial lipopolysaccharides. Immunochemistry 13, 813–818

6. Boll, J. M., Crofts, A. A., Peters, K., Cattoir, V., Vollmer, W., Davies, B. W., and Trent, M. S. (2016) A penicillin-binding protein inhibits selection of colistin-resistant, lipooligosaccharide-deficient Acinetobacter baumannii. Proc Natl Acad Sci U S A 113, E6228–E6237

7. Peng, D., Hong, W., Choudhury, B. P., Carlson, R. W., and Gu, X. X. (2005) Moraxella catarrhalis bacterium without endotoxin, a potential vaccine candidate. Infect Immun 73, 7569–7577

8. Steeghs, L., den Hartog, R., den Boer, A., Zomer, B., Roholl, P., and van der Ley, P. (1998) Meningitis bacterium is viable without endotoxin. Nature 392, 449–450

9. Zik, J. J., Yoon, S. H., Guan, Z., Stankeviciute Skidmore, G., Gudoor, R. R., Davies, K. M., Deutschbauer, A. M., Goodlett, D. R., Klein, E. A., and Ryan, K. R. (2022) Caulobacter lipid A is conditionally dispensable in the absence of fur and in the presence of anionic sphingolipids. Cell Rep 39, 110888

10. Smit, J., Kaltashov, I. A., Cotter, R. J., Vinogradov, E., Perry, M. B., Haider, H., and Qureshi, N. (2008) Structure of a novel lipid A obtained from the lipopolysaccharide of Caulobacter crescentus. Innate Immun 14, 25–37

11. Gomez-Munoz, A. (2004) Ceramide-1-phosphate: a novel regulator of cell activation. FEBS Lett 562, 5–10

12. Sugiura, M., Kono, K., Liu, H., Shimizugawa, T., Minekura, H., Spiegel, S., and Kohama, T. (2002) Ceramide kinase, a novel lipid kinase. Molecular cloning and functional characterization. J Biol Chem 277, 23294–23300

13. Ranjit, D. K., Moye, Z. D., Rocha, F. G., Ottenberg, G., Nichols, F. C., Kim, H. M., Walker, R., Gibson, F. C., 3rd, and Davey, M. E. (2022) Characterization of a bacterial kinase that phosphorylates dihydrosphingosine to form dhS1P. Microbiol Spectr 10, e0000222

14. De Siervo, A. J., and Homola, A. D. (1980) Analysis of Caulobacter crescentus lipids. J Bacteriol 143, 1215–1222

15. Don, A. S., and Rosen, H. (2008) A fluorescent plate reader assay for ceramide kinase. Anal Biochem 375, 265–271

16. Labesse, G., Douguet, D., Assairi, L., and Gilles, A. M. (2002) Diacylglyceride kinases, sphingosine kinases and NAD kinases: distant relatives of 6-phosphofructokinases. Trends Biochem Sci 27, 273–275

17. Lidome, E., Graf, C., Jaritz, M., Schanzer, A., Rovina, P., Nikolay, R., and Bornancin, F. (2008) A conserved cysteine motif essential for ceramide kinase function. Biochimie 90, 1560–1565

18. Graf, C., Klumpp, M., Habig, M., Rovina, P., Billich, A., Baumruker, T., Oberhauser, B., and Bornancin, F. (2008) Targeting ceramide metabolism with a potent and specific ceramide kinase inhibitor. Mol Pharmacol 74, 925–932

19. Han, G. S., O’Hara, L., Carman, G. M., and Siniossoglou, S. (2008) An unconventional diacylglycerol kinase that regulates phospholipid synthesis and nuclear membrane growth. J Biol Chem 283, 20433–20442

20. Bakali, H. M., Herman, M. D., Johnson, K. A., Kelly, A. A., Wieslander, A., Hallberg, B. M., and Nordlund, P. (2007) Crystal structure of YegS, a homologue to the mammalian diacylglycerol kinases, reveals a novel regulatory metal binding site. J Biol Chem 282, 19644–19652

21. Stankeviciute, G., Tang, P., Ashley, B., Chamberlain, J. D., Hansen, M. E. B., Coleman, A., D’Emilia, R., Fu, L., Mohan, E. C., Nguyen, H., Guan, Z., Campopiano, D. J., and Klein, E. A. (2022) Convergent evolution of bacterial ceramide synthesis. Nat Chem Biol 18, 305–312

22. Heaver, S. L., Le, H. H., Tang, P., Basle, A., Mirretta Barone, C., Vu, D. L., Waters, J. L., Marles-Wright, J., Johnson, E. L., Campopiano, D. J., and Ley, R. E. (2022) Characterization of inositol lipid metabolism in gut-associated Bacteroidetes. Nat Microbiol 7, 986–1000

23. Kawahara, K., Moll, H., Knirel, Y. A., Seydel, U., and Zähringer, U. (2000) Structural analysis of two glycosphingolipids from the lipopolysaccharide-lacking bacterium Sphingomonas capsulata. European Journal of Biochemistry 267, 1837–1846

24. Stankeviciute, G., Guan, Z., Goldfine, H., and Klein, E. A. (2019) Caulobacter crescentus adapts to phosphate starvation by synthesizing anionic glycoglycerolipids and a novel glycosphingolipid. mBio 10, e00107–00119

25. Kanzaki, H., Movila, A., Kayal, R., Napimoga, M. H., Egashira, K., Dewhirst, F., Sasaki, H., Howait, M., Al-Dharrab, A., Mira, A., Han, X., Taubman, M. A., Nichols, F. C., and Kawai, T. (2017) Phosphoglycerol dihydroceramide, a distinctive ceramide produced by Porphyromonas gingivalis, promotes RANKL-induced osteoclastogenesis by acting on non-muscle myosin II-A (Myh9), an osteoclast cell fusion regulatory factor. Biochim Biophys Acta Mol Cell Biol Lipids 1862, 452–462

26. Brown, E. M., Ke, X., Hitchcock, D., Jeanfavre, S., Avila-Pacheco, J., Nakata, T., Arthur, T. D., Fornelos, N., Heim, C., Franzosa, E. A., Watson, N., Huttenhower, C., Haiser, H. J., Dillow, G., Graham, D. B., Finlay, B. B., Kostic, A. D., Porter, J. A., Vlamakis, H., Clish, C. B., and Xavier, R. J. (2019) Bacteroides-derived sphingolipids are critical for maintaining intestinal homeostasis and symbiosis. Cell Host Microbe 25, 668–680 e667

27. Yu, N. Y., Wagner, J. R., Laird, M. R., Melli, G., Rey, S., Lo, R., Dao, P., Sahinalp, S. C., Ester, M., Foster, L. J., and Brinkman, F. S. (2010) PSORTb 3.0: improved protein subcellular localization prediction with refined localization subcategories and predictive capabilities for all prokaryotes. Bioinformatics 26, 1608–1615

28. Moye, Z. D., Valiuskyte, K., Dewhirst, F. E., Nichols, F. C., and Davey, M. E. (2016) Synthesis of sphingolipids impacts survival of Porphyromonas gingivalis and the presentation of surface polysaccharides. Front Microbiol 7, 1919

29. Rocha, F. G., Moye, Z. D., Ottenberg, G., Tang, P., Campopiano, D. J., Gibson, F. C., 3rd, and Davey, M. E. (2020) Porphyromonas gingivalis sphingolipid synthesis limits the host inflammatory response. J Dent Res 99, 568–576

30. Walsh, J. P., and Bell, R. M. (1986) sn-1,2-Diacylglycerol kinase of Escherichia coli. Mixed micellar analysis of the phospholipid cofactor requirement and divalent cation dependence. J Biol Chem 261, 6239–6247

31. Bligh, E. G., and Dyer, W. J. (1959) A rapid method of total lipid extraction and purification. Can J Biochem Physiol 37, 911–917

32. Goldfine, H., and Guan, Z. (2015) Lipidomic Analysis of Bacteria by Thin-Layer Chromatography and Liquid Chromatography/Mass Spectrometry. in Hydrocarbon and Lipid Microbiology Protocols (McGenity, T. J. ed.), Humana Press. pp 1–15

33. Guan, Z., Katzianer, D., Zhu, J., and Goldfine, H. (2014) Clostridium difficile contains plasmalogen species of phospholipids and glycolipids. Biochim Biophys Acta 1842, 1353–1359

34. Schindelin, J., Arganda-Carreras, I., Frise, E., Kaynig, V., Longair, M., Pietzsch, T., Preibisch, S., Rueden, C., Saalfeld, S., Schmid, B., Tinevez, J. Y., White, D. J., Hartenstein, V., Eliceiri, K., Tomancak, P., and Cardona, A. (2012) Fiji: an open-source platform for biological-image analysis. Nat Methods 9, 676–682

35. Altschul, S. F., Gish, W., Miller, W., Myers, E. W., and Lipman, D. J. (1990) Basic local alignment search tool. J Mol Biol 215, 403–410

36. Edgar, R. C. (2004) MUSCLE: a multiple sequence alignment method with reduced time and space complexity. BMC Bioinformatics 5, 113

37. Stamatakis, A. (2014) RAxML version 8: a tool for phylogenetic analysis and post-analysis of large phylogenies. Bioinformatics 30, 1312–1313

38. Miller, M. A., Pfeiffer, W., and Schwartz, T. (2010) Creating the CIPRES Science Gateway for inference of large phylogenetic trees. 2010 Gateway Computing Environments Workshop (GCE), 1–8

39. Yu, G. (2020) Using ggtree to visualize data on tree-like structures. Curr Protoc Bioinformatics 69, e96

40. Paradis, E., and Schliep, K. (2019) ape 5.0: an environment for modern phylogenetics and evolutionary analyses in R. Bioinformatics 35, 526–528

41. Wang, L. G., Lam, T. T., Xu, S., Dai, Z., Zhou, L., Feng, T., Guo, P., Dunn, C. W., Jones, B. R., Bradley, T., Zhu, H., Guan, Y., Jiang, Y., and Yu, G. (2020) Treeio: An R Package for phylogenetic tree input and output with richly annotated and associated data. Mol Biol Evol 37, 599–603

42. Wickham, H. (2016) ggplot2: Elegant Graphics for Data Analysis Springer-Verlag New York

